# Enhancing violacein production in an Antarctic bacterial strain: New opportunities for biomedical use

**DOI:** 10.64898/2026.11.16.766438

**Authors:** Hany Pazos-Espinosa, Eugenia Vila, Guillermo Valdomir, Martín Bacini, Alexandra Castro, Lucía Canclini, Diego Alem

**Affiliations:** Departamento de Genética, Instituto de Investigaciones Biológicas Clemente Estable, Montevideo, Uruguay; Departamento de Ingeniería Bioquímica y Bioprocesos, Facultad de Ingeniería, Universidad de la República, Montevideo, Uruguay; Laboratorio de Química Farmacéutica, Departamento de Química Orgánica, Facultad de Química, Universidad de la República, Montevideo, Uruguay; Laboratorio de Biología Molecular Vegetal, Instituto de Química Biológica, Facultad de Ciencias, Universidad de la República, Montevideo, Uruguay; Departamento de Medicina, Universidad Católica del Uruguay, Montevideo, Uruguay

**Keywords:** natural pigment, *Janthinobacterium* sp., Antarctica, Bioprocess optimization, violacein

## Abstract

Violacein is a valuable bioactive metabolite produced by diverse bacteria, including the Antarctic isolate *Janthinobacterium* sp. UV13. Despite its broad therapeutic potential, large-scale production in native producers is hindered by low yields and the concurrent synthesis of exopolysaccharides, which increases downstream processing complexity and costs. This study employed a sequential statistical approach to optimize culture parameters, enhancing violacein productivity while minimizing exopolysaccharides production in UV13. The methodology included carbon and nitrogen source screening, a Plackett–Burman design and a concentration–response optimization, followed by chemical inducers evaluation. Results demonstrated that sucrose was an effective carbon source (violacein yield: 32.75 ± 1.07 mg/L). Inorganic nitrogen significantly favored violacein synthesis (yield: 26 - 28 mg/L, *p* < 0.05) over organic alternatives which reduced violacein yields 3.4- to 4.2-fold. A single-factor central composite design of ammonium sulfate concentration yielded a formulation producing 75.8 mg/L of violacein with an exopolysaccharide titer of 0.91 mg/L. Furthermore, tryptophan was the better inducer, increasing violacein production 2.2-fold (*p* < 0.05) while simultaneously decreasing exopolysaccharides formation 4.5-fold. The new formulated medium significantly outperformed previously reported conditions, demonstrating that parameter manipulation unlocks UV13’s biosynthetic potential for scalable violacein production.

## 1. Introduction

Violacein is a violet secondary metabolite produced by diverse bacterial genera, including *Chromobacterium*, *Janthinobacterium*, *Rugamonas*, *Duganella*, *Massilia*, *Pseudoalteromonas*, and *Collimonas* [1–3]. This bisindole compound is synthesized via the *vioABCDE* operon, which catalyzes the oxidative condensation of two L-tryptophan molecules through a conserved enzymatic pathway [4–6]. While its primary biological role remains to be fully elucidated, studies report that this secondary metabolite confers resistance to environmental stress and is often regulated by quorum sensing mechanisms [7–9].

Beyond its ecological role, violacein has gained significant attention due to its broad-spectrum therapeutic potential, including immunomodulatory, antimicrobial, antiparasitic, and antitumoral properties [1,3]. Ongoing progress in understanding its mechanisms and toxicity have further highlighted its potential to combat multidrug-resistant pathogens, target cancer hallmarks, and provide neuroprotective effects against amyotrophic lateral sclerosis [10–12]. Consequently, optimizing the bioproduction of this versatile and promising metabolite has become a priority for ensuring its viability in industrial applications.

Strategies to increase violacein yields include optimizing culture conditions in native producers, as well as the heterologous expression of *vioABCDE* operon in industrial model microorganisms [13–17]. Evidence indicates that violacein production is highly sensitive to physicochemical parameters, such as temperature, pH, medium composition, agitation, feeding regimes, with these effects being strictly strain-dependent [17–19]. Despite these advancements, achieving economically viable industrial-scale production remains a challenge, requiring the design of scalable and cost-effective production strategies. Furthermore, since commercially available standards reach a maximum purity of approximately 85% [20], there is a need for integrated production and downstream purification strategies to obtain higher-purity preparations.

Our group has demonstrated violacein production from the Antarctic psychrotolerant strain *Janthinobacterium* sp. UV13 [21]. Previous bioreactor-scale studies identified that cultivation in commercial tryptic soy broth (TSB) supplemented with glucose optimized violacein production for UV13. This approach achieved a 4.8% recovery yield, a value higher than those reported for other natural producers, such as *Chromobacterium* sp. NIIST (0.8%; [22]) and *Janthinobacterium lividum* (4.5 %;[17]). However, Alem *et al*. [21] reported that these conditions also triggered the concurrent synthesis of notable amounts of exopolysaccharides (EPS) in UV13. In microbial pigment production processes, EPS is often an undesirable secondary outcome. Its production leads to a partitioning of nutrients toward non-target metabolites, while its accumulation alters the rheological properties of the medium, hindering broth mixing and aeration by reducing the overall volumetric oxygen mass transfer coefficient [23] and hampering downstream purification, thereby complicating and raising the target recovery cost [24]. Thus, the aim of this work was to optimize culture parameters using statistical methodologies to enhance violacein productivity in *Janthinobacterium* sp. UV13 while reducing EPS production. To this end, variables such as carbon and nitrogen sources, specific chemical inducers, aeration, and cultivation time were comprehensively investigated.

## 2. Results

### 2.1. Carbon and Nitrogen source screening

To evaluate the nutritional requirements for optimal cell growth and pigment production, different carbon and nitrogen sources were tested in a modified Davis medium [32] (Fig. 1). Regarding biomass production (Fig. 1a), sucrose supported the highest cell growth, reaching a dry cell weight (DCW) of 1.18 ± 0.05 g/L. In contrast, cultures supplemented with D-glucose or glycerol showed significantly lower biomass yields of 0.68 ± 0.10 and 0.70 ± 0.10 g/L, respectively. Methanol and D-xylose failed to support substantial growth of strain UV13 under the tested conditions. Consistent with these results, sucrose was also the most effective carbon source for violacein production, yielding 32.75 ± 1.07 mg/L (Fig. 1b). Conversely, violacein synthesis was markedly reduced in cultures supplemented with D-glucose or glycerol. These differences were also visually apparent from the intense purple coloration of sucrose-supplemented cultures (Fig. 1c), further confirming the superior performance of sucrose in supporting both biomass accumulation and pigment production.

**Figure 1.**
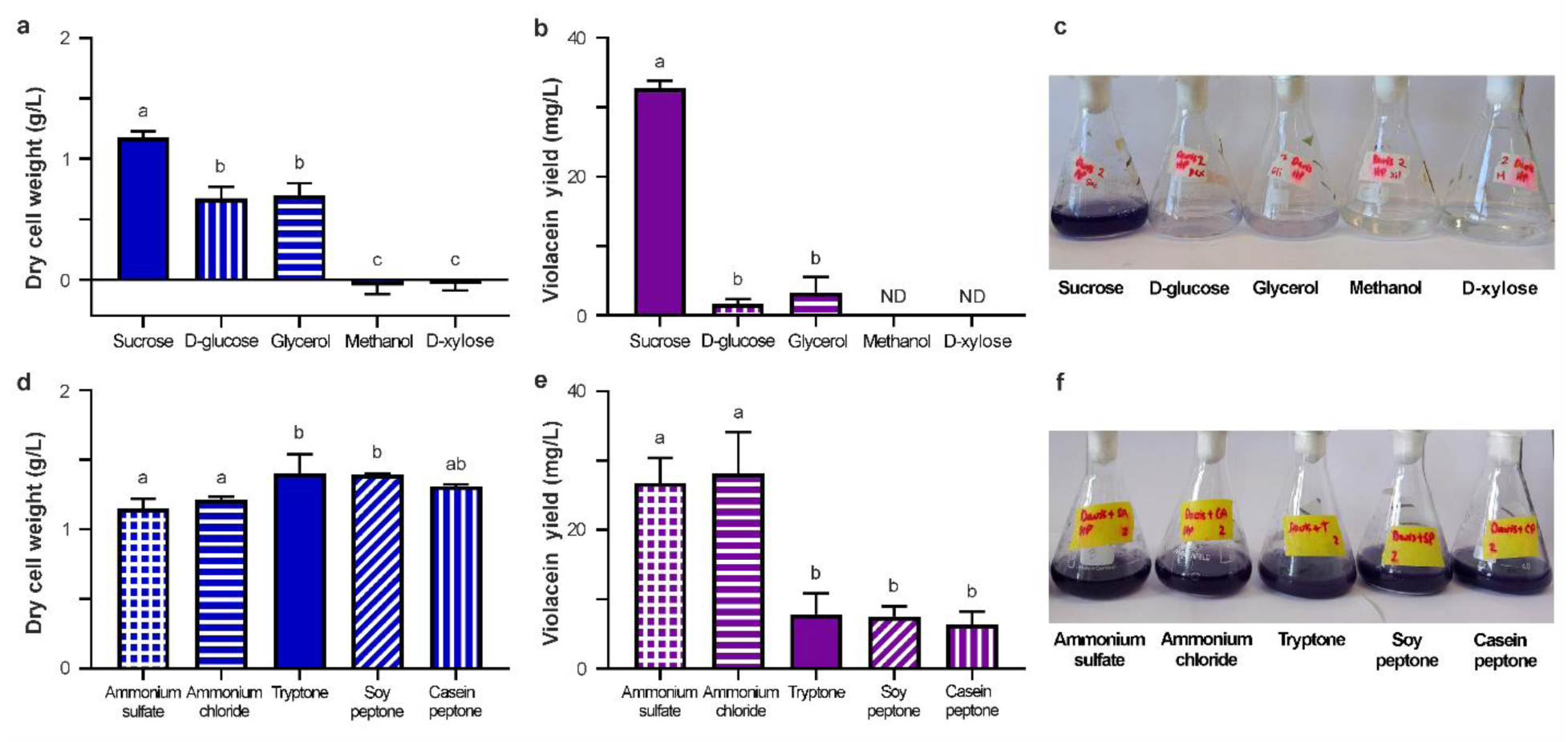
Screening of carbon and nitrogen sources on *Janthinobacterium* sp. UV13 grown in a modified Davis synthetic medium after 72 h of incubation. (a) Dry cell weight and (b) Violacein yield of the strain cultivated on different carbon sources. (d) Dry cell weight and (e) Violacein yield of the strain cultivated on different nitrogen sources. (c,f) Visual appearance of culture flasks at 72 h endpoint. Graphs show the mean ± standard deviation of biological triplicates. Different lowercase letters indicate statistically significant differences between conditions (*p* < 0.05) determined by one-way ANOVA with Tukey’s post hoc test for multiple comparisons. ND: not detected.

Regarding the evaluation of nitrogen sources, distinct trends were observed between biomass and pigment accumulation. For biomass, the highest DCW levels of 1.40 ± 0.14 and 1.40 ± 0.01 g/L were achieved using tryptone and soy peptone, respectively (Fig. 1d). Notably, an apparent increase in medium viscosity was observed during cultivation with these organic components, although it was not quantitatively measured. Conversely, cultures utilizing tryptone, soy peptone, or casein peptone exhibited a significant decrease in violacein yields, with reductions ranging from 3.4- to 4.2-fold. Instead, the inorganic nitrogen sources ammonium sulfate and ammonium chloride significantly outperformed the other components, driving the highest violacein accumulation at 26.79 ± 3.59 and 28.18 ± 5.94 mg/L, respectively (Fig. 1e). Despite these quantitative differences, all tested sources supported growth and violacein production, as indicated by the purple color of all culture flasks in Figure 1f.

### 2.2. Plackett–Burman design

To identify the nutritional factors that enhance violacein production and biomass accumulation, a Plackett–Burman design was implemented at a 72 h endpoint. The eight nutritional components tested within the modified Davis medium formulation were detailed in the Materials and Methods section. The results for each monitored response (dry cell weight, specific and volumetric violacein yields, and specific and volumetric EPS yields) are summarized in Table 1, and the culture visual performance is presented in Supplementary Figure S1.

**Table 1.**
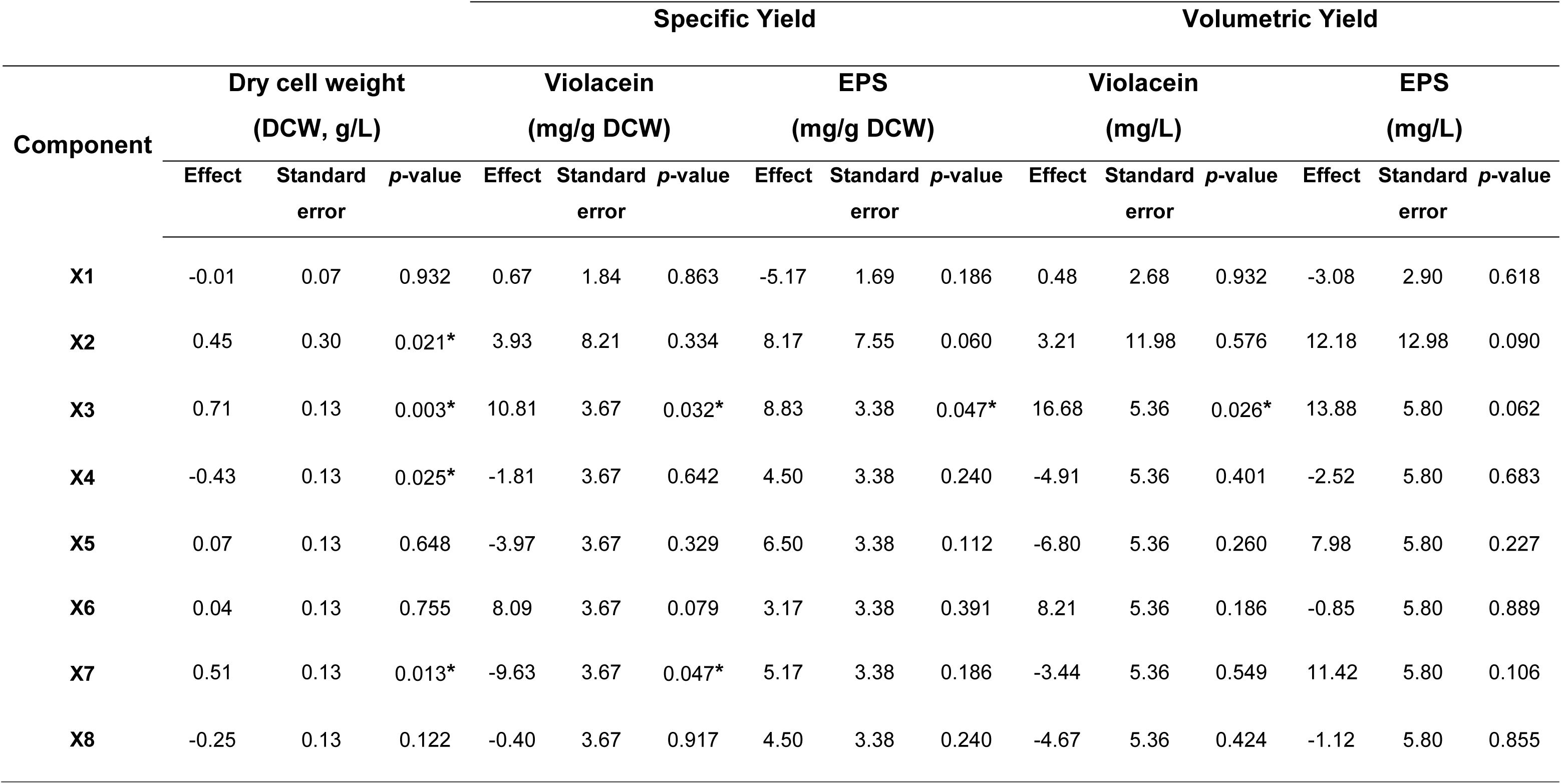
Statistical evaluation of a modified Davis synthetic medium components using a Plackett–Burman design: Main effects and significance levels for Dry cell weight, Specific and Volumetric violacein yield, Specific and Volumetric EPS yield at 72 h endpoint.

|  |  |  |  | **Specific Yield** |  |  |  |  |  | **Volumetric Yield** |  |  |  |  |  |
|---|---|---|---|---|---|---|---|---|---|---|---|---|---|---|---|
| **Component** | **Dry cell weight<br>(DCW, g/L)** |  |  | **Violacein<br>(mg/g DCW)** |  |  | **EPS<br>(mg/g DCW)** |  |  | **Violacein<br>(mg/L)** |  |  | **EPS<br>(mg/L)** |  |  |
|  | **Effect** | **Standard error** | ***p*-value** | **Effect** | **Standard error** | ***p*-value** | **Effect** | **Standard error** | ***p*-value** | **Effect** | **Standard error** | ***p*-value** | **Effect** | **Standard error** | ***p*-value** |
| **X1** | -0.01 | 0.07 | 0.932 | 0.67 | 1.84 | 0.863 | -5.17 | 1.69 | 0.186 | 0.48 | 2.68 | 0.932 | -3.08 | 2.90 | 0.618 |
| **X2** | 0.45 | 0.30 | 0.021* | 3.93 | 8.21 | 0.334 | 8.17 | 7.55 | 0.060 | 3.21 | 11.98 | 0.576 | 12.18 | 12.98 | 0.090 |
| **X3** | 0.71 | 0.13 | 0.003* | 10.81 | 3.67 | 0.032* | 8.83 | 3.38 | 0.047* | 16.68 | 5.36 | 0.026* | 13.88 | 5.80 | 0.062 |
| **X4** | -0.43 | 0.13 | 0.025* | -1.81 | 3.67 | 0.642 | 4.50 | 3.38 | 0.240 | -4.91 | 5.36 | 0.401 | -2.52 | 5.80 | 0.683 |
| **X5** | 0.07 | 0.13 | 0.648 | -3.97 | 3.67 | 0.329 | 6.50 | 3.38 | 0.112 | -6.80 | 5.36 | 0.260 | 7.98 | 5.80 | 0.227 |
| **X6** | 0.04 | 0.13 | 0.755 | 8.09 | 3.67 | 0.079 | 3.17 | 3.38 | 0.391 | 8.21 | 5.36 | 0.186 | -0.85 | 5.80 | 0.889 |
| **X7** | 0.51 | 0.13 | 0.013* | -9.63 | 3.67 | 0.047* | 5.17 | 3.38 | 0.186 | -3.44 | 5.36 | 0.549 | 11.42 | 5.80 | 0.106 |
| **X8** | -0.25 | 0.13 | 0.122 | -0.40 | 3.67 | 0.917 | 4.50 | 3.38 | 0.240 | -4.67 | 5.36 | 0.424 | -1.12 | 5.80 | 0.855 |

Statistical analysis revealed that violacein production was significantly promoted by X3, whereas X7 exhibited a marked negative effect on specific violacein yield (*p* = 0.032). Regarding biomass accumulation, four of the eight evaluated components exerted significant influences. Specifically, increasing concentrations of X2, X3, and X7 showed positive effects and significantly enhanced cell growth, whereas X4 acted as an inhibitory factor within the tested concentration range, showing a significant negative effect on biomass accumulation. For EPS production, only X3 stood out as a statistically significant component driving specific yield, exerting a positive effect.

Taken together, as X3 acts as a crucial master regulator for both growth and metabolite upregulation, it was selected for further optimization study. The analysis also indicated that factors like X4 and X7 introduce critical trade-offs between biomass accumulation and targeted pigment yields; thus, they were excluded from subsequent formulation. Finally, X2 was fixed at its high level, and components with no significant effect on these monitored responses, were kept at the central level.

### 2.3. Growth medium optimization for violacein production

The effects of ammonium sulfate concentration on violacein production, biomass and EPS accumulation were evaluated using a Central Composite Design (CCD). As shown in Figure 2, the quadratic models adequately fitted each response within the ammonium sulfate range tested. Conversely, dry cell weight and EPS yield, displayed parabolic profiles. A maximum accumulation of 1.63 g/L of DCW is expected at 1.25 g/L of ammonium sulfate (Fig. 2a). Violacein yield decreased over the range tested, with a non-significant quadratic term (*p* = 0.1117), reaching its maximum expected production of 79.59 mg/L at the lowest tested concentration of 0.22 g/L of ammonium sulfate (Fig. 2b). Regarding EPS yield, its lowest production zone was observed at the lowest tested concentration of 0.22 g/L ammonium sulfate (Fig. 2c). Thus, this lower boundary represented the optimal criterion to maximize violacein yield while balancing biomass and minimizing EPS accumulation. To verify model accuracy, a validation experiment was conducted at the optimal 0.22 g/L ammonium sulfate concentration. The assay yielded 75.84 mg/L of violacein, 1.69 g/L of DCW, and 0.91 mg/L of EPS, exhibiting relative errors of 4.94%, 8.46%, and 63.00% against predicted values, respectively. The regression parameters for the fitted models are summarized in Fig. 2d. All models were statistically significant (*p* < 0.005) with high coefficients of determination (*R*^2^ > 89 %), accounting for most of the experimental variability. Additionally, the non-significant lack of fit (*p* > 0.05) across all responses confirmed the mathematical reliability and accuracy of the quadratic models to describe the system.

**Figure 2.**
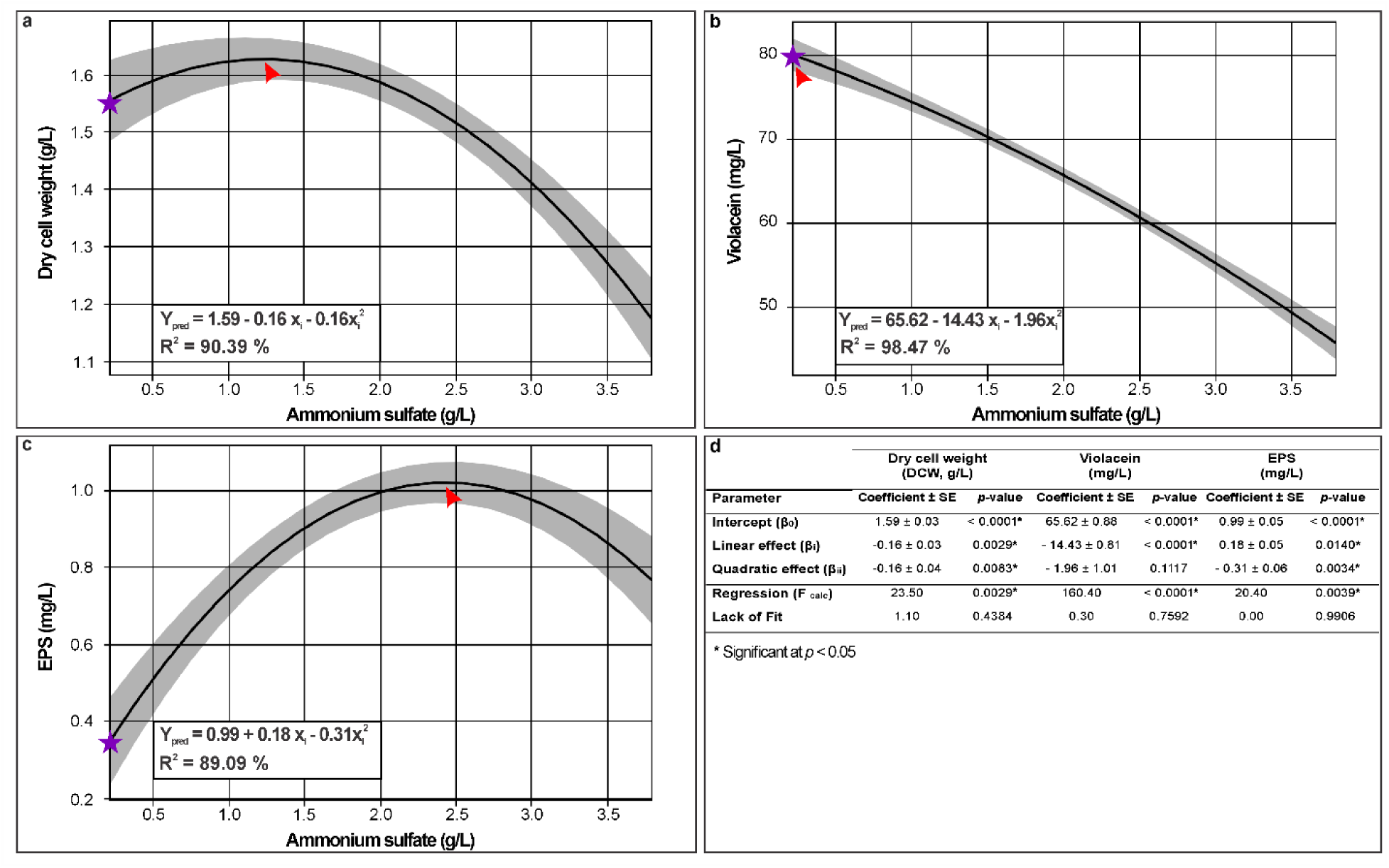
Optimization profiles as a function of ammonium sulfate concentration. (a) Dry cell weight, (b) Violacein yield, (c) EPS yield and d) Statistical data analysis. The operational ranges for the independent variable correspond to the experimental domain detailed in Section 3.5. The purple star marks the selected optimal criterion, while the red arrowheads indicate the points of maximum expected production for each response.

### 2.4. Chemical Inducers to Enhance Violacein Production

To identify effective chemical inducers for maximizing pigment yields, the effects of formic acid, ampicillin, and tryptophan were compared at three doses each (Fig. 3). The addiction of all inductors generated a reduction in DCW except formic acid as the low concentration (Fig. 3a). Culture flasks with an evident violet increase with tryptophane were observed (Fig. 3b), which was identified as the only candidate that successfully upregulated violacein production in UV13 regarding control conditions. Specific violacein yield increased by at least 175% (p < 0.0001, Fig. 3c), while volumetric yield increased by at least 50% (p < 0.001, Fig. 3d) compared to the control. Within the tested tryptophan range, violacein production reached a plateau between 0.3 and 0.6 mg/mL. Increasing the dosage beyond 0.3 mg/mL did not further suppress EPS production below baseline levels. Conversely, formic acid and ampicillin acted as opposing metabolic triggers, driving production entirely away from the target pigment. Both supplements significantly suppressed specific and volumetric violacein yields in a dose-dependent manner. At their highest tested doses, ampicillin and formic acid caused a complete inhibition or a visible reduction of violacein production, respectively (p < 0.0001, Fig. 3c, d). Concurrently, they stimulated a dose-dependent overproduction of EPS, with an EPS spike when ampicillin was supplied at 25 µg/mL (p < 0.0001, Fig. 3c, d).

**Figure 3.**
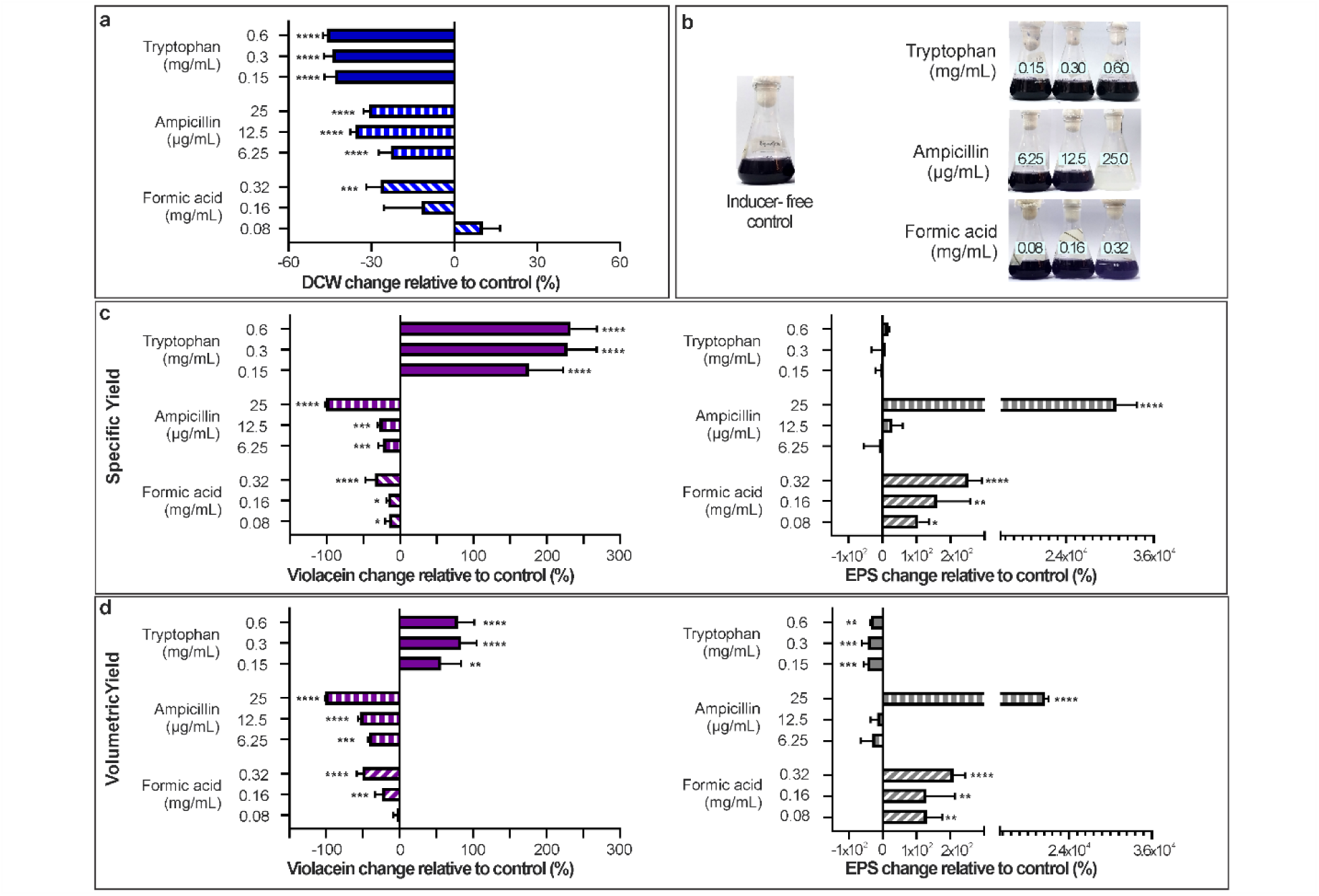
Evaluation of inducer-driven metabolic effects on violacein, biomass, and EPS yields. (a) Dry cell weight (DCW) changes relative to the inducer-free control. (b) Visual appearance of culture flasks compared to the inducer-free control. (c) Specific yields and (d) volumetric yields showcasing changes in violacein (purple bars) and EPS (gray bars) relative to the inducer-free control. Data represent the mean ± standard deviation of biological triplicates. Asterisks indicate statistically significant differences relative to the control condition (\**p* < 0.05, \*\**p* < 0.01, \*\*\**p* < 0.001, \*\*\*\**p* < 0.0001) determined by one-way ANOVA with Dunnett’s post hoc test for multiple comparisons.

### 2.5. Effect of aeration and cultivation time

To evaluate the influence of operational conditions on *Janthinobacterium* sp. UV13 grown in the optimized HPE-VP medium, the effects of aeration and cultivation time were investigated (Fig. 4). Increasing aeration by using a 1:5 medium-to-flask volume ratio significantly enhanced biomass accumulation (Fig. 4a) while maintaining specific violacein yields without increasing EPS production (Fig. 4b). Consequently, volumetric violacein yield reached 168.6 mg/L, compared with 120.9 mg/L under the control 1:2 ratio after 72 h of cultivation (Fig. 4c). This condition also resulted in the highest volumetric violacein productivity, reaching 2.34 mg/L·h compared with 1.68 mg/L·h under the control condition. In contrast, extending cultivation to 144 h under the 1:2 ratio resulted in a volumetric violacein yield of 160.0 mg/L, but reduced productivity to its lowest value (1.11 mg/L·h). Moreover, prolonged cultivation significantly increased volumetric EPS production by 2.19-fold, reaching 2.46 mg/L (Fig. 4c).

**Figure 4.**
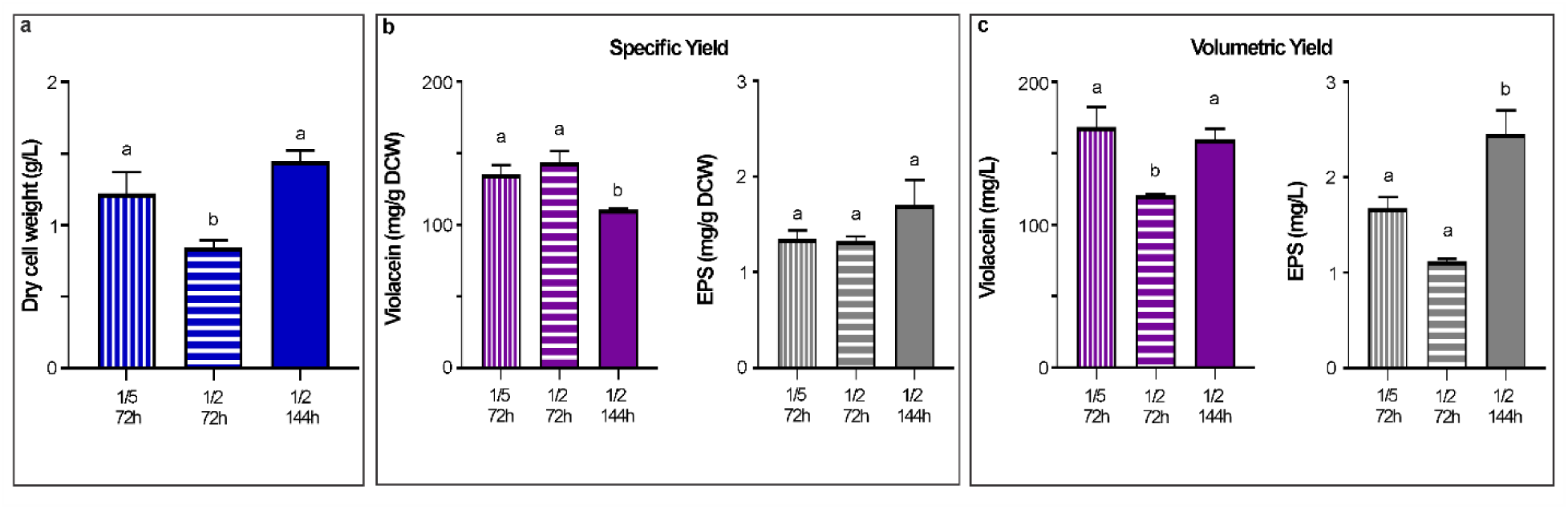
Effects of aeration and cultivation time on *Janthinobacterium* sp. UV13 grown in the optimized HPE-VP medium. (a) Dry cell weight (DCW); (b) Specific yields of violacein (purple bars) and EPS (gray bars); (c) Volumetric yields of violacein (purple bars) and EPS (gray bars). Data represent the mean ± standard deviation of biological triplicates. Different lowercase letters above the bars indicate statistically significant differences between treatments according to one-way ANOVA with Tukey’s post hoc test for multiple comparisons (*p* < 0.05).

### 2.6. Comparative evaluation with Baseline Medium

To evaluate the performance of the optimized HPE-VP medium against baseline medium (TSB+Glucose) reported by Alem *et al.* [21], comparative cultures were analyzed (Fig. 5). Regarding cell growth, HPE-VP allowed a biomass accumulation of 1.2 g/L, which was 3.7-fold lower than that prompted by TSB+Glucose (*p* < 0.0001) (Fig 5a). However, using HPE-VP, UV13 achieved a significantly higher specific violacein and volumetric yields of 135.6 mg/g DCW and 168.6 mg/L, respectively (Fig. 5b, c). TSB+Glucose resulted in a drastically lower pigment yield, exhibiting 17.4 mg/g DCW (*p* < 0.0022) and 76.2 mg/L (*p* < 0.0063). These results are consistent with the increased transcript of *Vio*A and *Vio*D within the violacein biosynthetic pathway in the optimized medium relative to the baseline (Fig. 5d). Moreover, both specific and volumetric EPS yields were minimized in HPE-VP, while they were significantly favored in TSB+Glucose (*p* < 0.05 and *p* < 0.01, respectively) (Fig 5b, c). The reversed-phase chromatogram of the butanolic extract, recovered in acetonitrile:water (1:1), showed a 575nm profile with three peaks under both conditions (Supplementary Figure S2). The HPE-VP sample yielded a higher relative violacein purity (91.33% of the total peak area) compared to the TSB+Glucose sample (86.58%), resulting in purified violacein yields of 0.38 and 0.25 mg/mL of culture, respectively. The retention times for proviolacein, violacein, and deoxyviolacein were 6.2, 9.1 and 10.6 min, respectively. These retention times and their UV–Vis spectra were consistent with those reported by Lee *et al*. [25]. Violacein identity was confirmed via MS ions at m/z 344 [M+H]⁺ and m/z 342 [M−H]⁻ in ESI⁻. Its MS/MS fragmentation patterns were similar to Calemi *et al*. [26] reports. For deoxyviolacein and proviolacein, final identification was confirmed by the observation of their pseudomolecular ions (m/z 328 [M+H]⁺ in ESI⁺ and m/z 326 [M−H]⁻ in ESI⁻) and analysis of their MS/MS fragmentation patterns, with several fragments tentatively assigned and found to be consistent with their corresponding structures.

**Figure 5.**
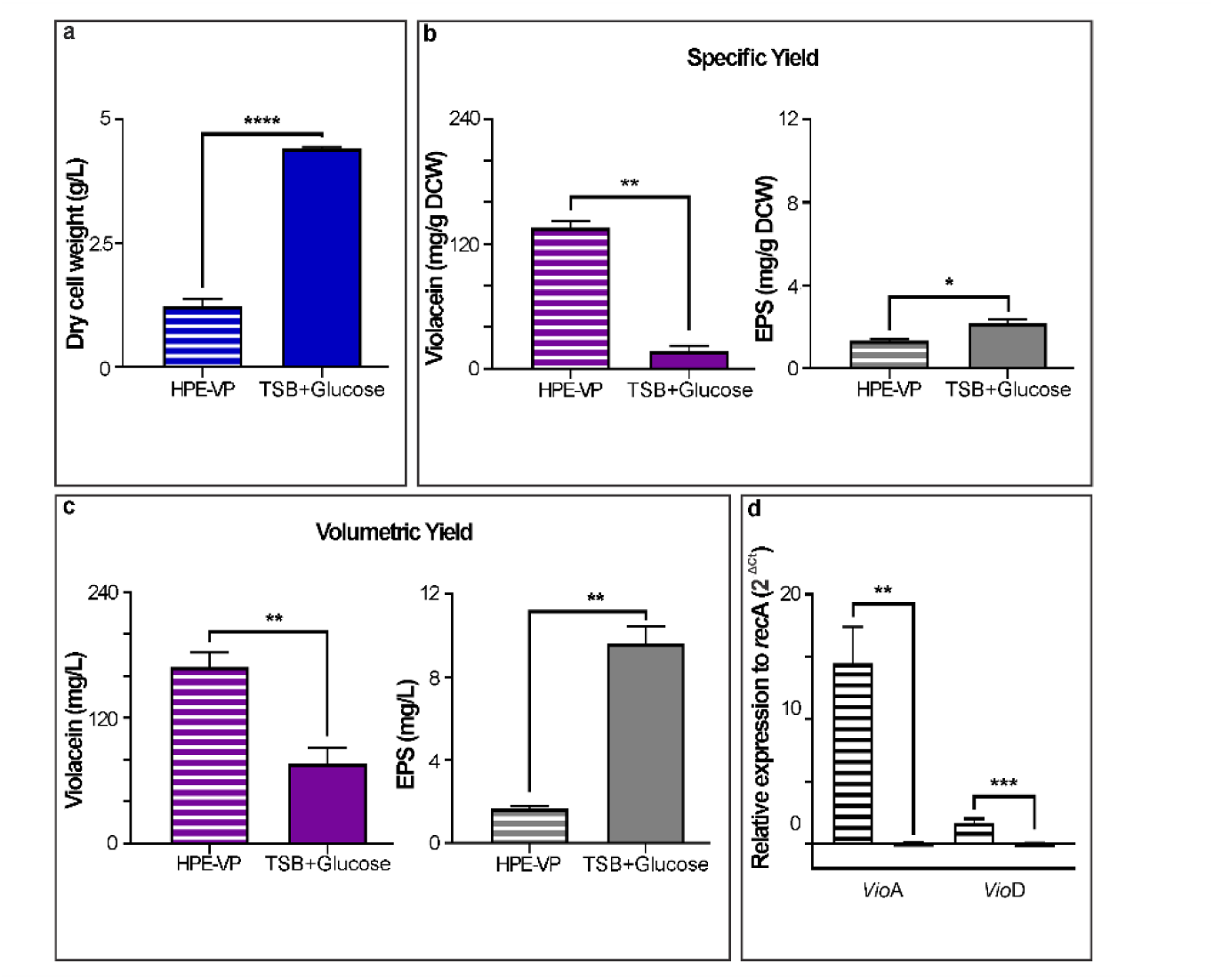
Comparative evaluation of the optimized HPE-VP medium against baseline medium on *Janthinobacterium* sp. UV13. (a) Dry cell weight (DCW); (b) Specific yields of violacein (purple bars) and EPS (gray bars); (c) Volumetric yields of violacein (purple bars) and EPS (gray bars). (d) Relative expression of *vio*A and *vio*D between both media conditions. Data represent the mean ± standard deviation of biological triplicates (striped bars: HPE-VP medium; solid bars: TSB+Glucose medium). Asterisks indicate statistically significant differences determined by t-test (\**p* < 0.05, \*\**p* < 0.01, \*\*\**p* < 0.001,\*\*\*\**p* < 0.0001).

## 3. Discussion

Violacein production is well documented across different bacterial genera; however, its synthesis remains highly sensitive to strain dependent physicochemical and nutritional parameters [1]. *Janthinobacterium* sp. UV13 was previously studied by Alem *et al.* [21] under different growth conditions in the commercial complex medium TSB, where limitations associated with excessive EPS production were identified. The present work is the first to evaluate a chemically defined medium formulation for this strain, screening different carbon and nitrogen sources, analyzing medium components, and adding inducers to enhance violacein production while seeking to avoid triggering excessive EPS accumulation.

Among the carbon sources evaluated, sucrose supported the best growth in this chemically defined medium formulation. Although glucose and glycerol effectively sustain this strain when supplemented into TSB [21], their concentration of 5.1 g/L on our synthetic medium limited overall growth. In synthetic media that have lower protective nutritional complexity of the TSB matrix, excessive amounts of these simple carbon substrates could induce substrate inhibition or disrupt the carbon to nitrogen balance, thereby reducing biomass production [27]. Within this scenario, sucrose provides metabolic advantages because this disaccharide must first be cleaved into its monosaccharide subunits prior to utilization. This controlled processing allows for a slower, steady metabolic feeding rate that avoids overflow metabolism [28]. Regarding nitrogen, inorganic ammonium salts efficiently channeled the metabolic flux toward maintaining active, pigment-producing cellular biomass that drives a better volumetric violacein yield in our strain. A similar metabolic preference for inorganic nitrogen sources over organic ones has been documented by Gharat & Singhal [29] for *Chromobacterium violaceum* MTCC2656. In UV13, complex organic nitrogen shifts the bacterial metabolism toward cell division and also triggers EPS production, which some authors propose operates as a carbon-sink mechanism [27, 30]. This shift toward EPS synthesis highlights a clear operational disadvantage of complex matrices for UV13 bioprocessing.

Optimizing the medium formulation is a crucial strategy in metabolic engineering to enhance violacein yields and improve bioprocess effectiveness. In this study, a Plackett-Burman analysis successfully screened the medium components, identifying ammonium sulfate as the critical factor for formulation. Subsequent optimization using a central composite design validated predictive models for both biomass and pigment production. Specifically, our model demonstrated a preference for restricted nitrogen levels to enhance violacein synthesis in our strain, since nitrogen richness shifts metabolic flux toward rapid biomass accumulation at the expense of pigment yield [31]. This behavior is consistent with other *Janthinobacterium* strains, which have also shown higher pigment production in minimal or nutrient-limited media [32].

Beyond the components optimized by statistical methodologies, the addition of chemical inducers was evaluated to further maximize violacein production. While ampicillin and formic acid have been reported to enhance yields in other natural producers, such as *J. lividum* 1522 strain ATCC 12473 [17] and *Chromobacterium violaceum* ATCC 12472 [9], respectively; they proved ineffective for violacein production in UV13. Instead, they appeared to trigger survival mechanisms, shifting the metabolic flux toward significant EPS overproduction at the expense of both biomass and pigment accumulation. This phenotypic decoupling suggests that violacein biosynthesis and EPS-mediated stress responses in UV13 may operate through independent regulatory mechanisms uncoupled from classical quorum sensing models [33–35].

In contrast, tryptophan supplementation stands as the optimal strategy to boost pigment production achieving a substantial increase of over 50% relative to the free-inducer control. Although our previous results reported that tryptophan supplementation failed to stimulate violacein yield when using a nutrient-rich commercial medium like TSB [21], in this new design, our findings indicate that it plays a critical role in nutrient-limited nutrient medium formulation. D*e novo* tryptophan synthesis imposes a high energetic and metabolic cost on the cell [36–37]. Because it serves as the direct metabolic precursor for the violacein biosynthetic pathway, supplying it exogenously likely bypasses this metabolic bottleneck. Consequently, this metabolic relief may shift the metabolic flux toward secondary metabolism driving violacein production without exacerbating EPS accumulation [38].

In addition to medium formulation, other operational parameters such as aeration and cultivation time, were preliminarily screened. Our results showed that the highest volumetric productivity of violacein occurred under a lower medium to air volume ratio, where enhanced dissolved oxygen and mass transfer boosted pigment production. This aligns with proposals by Durán *et al*. [1], where highlighted that higher oxygen transfer rate (OTR) supports effectively the strict energetic demands of tryptophan assimilation and enzymatic violacein synthesis. However, the metabolic dependency on oxygen availability remains a controversial and strain-dependent parameter within the genera. While a positive correlation with aeration has similarly been reported for *J. lividum* [39]; Frediansyah *et al*. [32] reported that violacein production in *Janthinobacterium agaricidamnosum* DSM 9628 and *J. lividum* DSM 1552 was higher under static conditions than under agitation across different synthetic and complex media. Therefore, fine tuning of OTR under our formulation, could be a fundamental manipulation strategy and a key criterion to consider during future scale-up for the UV13 isolate.

Under our optimal flasks culture conditions, the new formulation reached a greater than 2-fold increase in crude violacein yield over the baseline medium. This improved pigment productivity may have been associated with both the custom HPE-VP formulation, which promoted a 4.5-fold reduction in EPS accumulation, and the transcriptional upregulation of genes involved in the violacein biosynthetic pathway. EPS accumulation alters the rheological properties of the medium, hindering broth mixing and aeration by reducing the overall volumetric oxygen mass transfer coefficient [23]. The higher transcript levels of vioA and vioD are consistent with the increased violacein production observed in HPE-VP. VioA catalyzes the initial oxidation of L-tryptophan, whereas VioD participates in a later hydroxylation step leading to violacein formation [40]. Previous studies have shown that VioD expression can influence the relative production of violacein and deoxyviolacein [41]. However, the contribution of these transcriptional changes to violacein production in UV13 remains to be determined.

Furthermore, this optimization also outperformed previous bioreactor benchmarks for the UV13 isolate (77 mg/L of crude violacein, 4.8% pure pigment recovery yield) and also, some other natural producers within the genus *Janthinobacterium* at the same shake-flask scale, such as *J. agaricidamnosum* DSM 9628 and *J. lividum* DSM 1552 (9.8 mg/L and 10.89 mg/L, respectively [32]. While *J. lividum* XT1 achieved a higher yield of 3.50 g/L, its growth required a highly enriched medium [18]. Furthermore, our formulation places UV13 ahead of other violacein producing genera, including *Duganella violaceinigra* (18.6 mg/L) [42], *Chromobacterium* sp. NIIST (12.8 mg/L) [22], and the Antarctic *Iodobacter* sp. 7MAnt (1.1 mg/L) [19]. To further exploit this optimized formulation tailored for enhanced pigment production, evaluating parameters such as inoculum size and fed-batch operational modes represents a promising next step.

By significantly outperforming both our lab’s baseline and some other natural producers within the genus *Janthinobacterium*, this novel chemically defined formulation demonstrates that precise parameter tuning successfully unlocks UV13’s biosynthetic potential, providing a robust framework for economically viable violacein bioprocess scale-up.

## 4. Materials and Methods

### 4.1. Reagents, Bacterial strain and Culture conditions

Organic solvents were purchased from Dorwil. Tryptic soy broth (TSB), agar and tryptone were obtained from Oxoid. Tryptophan, magnesium sulfate, D-glucose, sucrose, glycerol, xylose, ammonium sulfate, ammonium chloride, soy peptone, casein peptone and HEPES were acquired from Merck-Millipore-Sigma. Sulfuric acid (95-98%) and formic acid (98-100%) were supplied by Cicarelli and Supelco, respectively. Dipotassium phosphate, monopotassium phosphate and sodium citrate were purchased from JT Baker. L-methionine and ampicillin were obtained from Applichem and BioChemica, respectively. The bacterial strain used was *Janthinobacterium* sp. UV13 kindly provided by Dr. Susana Castro-Sowinski (University of the Republic, Uruguay). For all experiments, inocula were prepared from single colonies and cultured in TSB (3 g/L) for 24 h at 20°C and 200 rpm. Unless stated otherwise, cultures were inoculated at an initial OD_660_ of 0.05 in 250 mL flasks containing 125 mL of medium, and grown at 20°C and 200 rpm for 72 h.

### 4.2. Analytical methods for biomass and metabolites determination

Cell growth was determined gravimetrically and expressed as dry cell weight per liter. Briefly, 5 mL aliquots of fermented broth were filtered through 0.22 μm membrane filters (Whatman, ME24/21ST) and dried at 60°C until a constant weight. All measurements were performed in triplicate, and a medium-only control was included.

For quantification of crude violacein, an ethanolic extract was obtained from the biomass under each tested culture condition. Briefly, 20 mL aliquots of fermented broth were centrifuged at 10,000 × g for 30 min and cell pellets were washed with 1X Phosphate-buffered saline (PBS). The biomass was lyophilized, and samples of known DCW were extracted with ethanol until the pellets were completely bleached. Violacein concentration was determined spectrophotometrically by measuring absorbance at 575 nm using a MultiSkan Sky plate reader (Thermo Scientific, USA). A violacein standard purified by Alem *et al.* [21] was used to generate a calibration curve in ethanol over the range 0 - 0.13 g/L, the corresponding linear regression equation was applied for quantification.

Exopolysaccharides (EPS) were quantified using the sulfuric acid-UV method described by Albalasmeh *et al*. [43] with modifications. Briefly, 5 mL aliquots of fermented broth were centrifuged and the cell pellets were washed with 1X PBS as described above. The biomass was then resuspended with 500 µL of 1X PBS, and 5 mL of cold isopropanol was added under agitation to precipitate the EPS. The precipitate was recovered by centrifugation, washed with ethanol and lyophilized. Samples of known DCW were resuspended with an aqueous sulfuric acid solution 0.05%. Subsequently, 200 µL of each sample was quickly mixed with 600 µL of concentrated sulfuric acid and incubated on ice for 1 h. Total carbohydrate concentration was determined spectrophotometrically by measuring absorbance at 315 nm using a ClarioStar Plus plate reader (BMG Labtech). A standard calibration curve prepared with glucose over the range 0 - 0.08 g/L was used, and quantification was based on the resulting linear regression equation.

### 4.3. Carbon and Nitrogen source screening

To evaluate the effect of medium composition on the growth and violacein production of *Janthinobacterium* sp. UV13 cultures were grown in a synthetic medium based on a modified Davis formulation previously applied to other violacein-producing strains [32]. The medium contained: 7.0 g/L dipotassium phosphate, 2.0 g/L monopotassium phosphate, 1.0 g/L ammonium sulfate, 0.5 g/L sodium citrate, 0.1 g/L magnesium sulfate, 0.56 g/L HEPES, 1 mg/L L-methionine and 1 mL/L trace metal mix (2.86 g H₃BO₃, 1.81 g MnCl₂·4H₂O, 0.222 g ZnSO₄·7H₂O, 0.390 mg Na₂MoO₄·2H₂O, 79 mg CuSO₄·5H₂O, 49 mg Co(NO₃)₂·6H₂O); initial pH was adjusted to 7.0. The carbon sources D-glucose, sucrose, glycerol, xylose, and methanol were screened at a final concentration of 5.1 g/L. The organic or inorganic nitrogen sources ammonium sulfate, ammonium chloride, tryptone, soy peptone, and casein peptone were also evaluated at an equivalent free nitrogen content. Cultivations were conducted under the standard incubation parameters described above, but adapted to a working volume of 63 mL in 125 mL flasks. All treatments were carried out in triplicate. Following incubation, cells were harvested as described above for subsequent analyses.

### 4.4. Identifying key nutrients using a Plackett–Burman design

To screen the impact of the synthetic medium components on the growth and violacein production of *Janthinobacterium* sp. UV13, a Plackett–Burman fractional factorial design was applied. The components and their tested coded levels [high (1), low (−1) and center (0)] are listed in Table 2. The experimental matrix comprised 12 factorial runs and three center points, all supplemented with 5.1 g/L sucrose and adjusted to pH 7.0. The Plackett–Burman design matrix for the eight components studied, including three center points, is provided in Supplementary Table S1. Cultures were performed as described above, and cells were harvested for all analysis.

**Table 2.** Concentration levels of the synthetic medium components evaluated in the Plackett–Burman fractional factorial design.

| **CODE** | **Components** | **Low level (-1)** | **Central level (0)** | **High level (1)** |
|---|---|---|---|---|
| **X1** | Dipotassium phosphate | 0 g/L | 3.5 g/L | 7.0 g/L |
| **X2** | Monopotassium phosphate | 0 g/L | 1 g/L | 2.0 g/L |
| **X3** | Ammonium sulfate | 0 g/L | 0.5 g/L | 1.0 g/L |
| **X4** | Sodium citrate | 0 g/L | 0.25 g/L | 0.5 g/L |
| **X5** | Magnesium sulfate | 0 g/L | 0.05 g/L | 0.1 g/L |
| **X6** | HEPES | 0 g/L | 0.28 g/L | 0.56 g/L |
| **X7** | L-methionine | 0 mg/L | 0.5 mg/L | 1 mg/L |
| **X8** | Trace metal mix | 0 mL/L | 0.5 mL/L | 1 mL/L |

### 4.5. Single-factor optimization of ammonium sulfate concentration

Based on the Plackett–Burman results, ammonium sulfate was selected for optimization using a Central Composite Design to maximize violacein production while minimizing EPS formation. The nutrient was tested at different coded levels, and the experimental matrix comprised four factorial runs and four center points. The coded design matrix and the responses measured are presented in Table 3. All runs were supplemented with 5.1 g/L sucrose and adjusted to pH 7.0. Cultures were performed as described above, and cells were harvested for all analyses. The response was modeled using a second-order polynomial equation (Eq. 1):

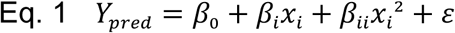

**Table 3.** Experimental matrix of the single-factor optimization for ammonium sulfate.

|  | **RUNS** |  |  |  |  |  |  |  |
|---|---|---|---|---|---|---|---|---|
| **Ammonium sulfate** | **Flask 1** | **Flask 2** | **Flask 3** | **Flask 4** | **Flask 5** | **Flask 6** | **Flask 7** | **Flask 8** |
| Coded levels | -1 | 1 | -1.19 | +1.19 | 0 | 0 | 0 | 0 |
| Concentration range (g/L) | 0.50 | 3.50 | 0.22 | 3.78 | 2.00 | 2.00 | 2.00 | 2.00 |

where:

Y_pred_: is the predicted response variable, β_0_: is the intercept coefficient, β_i_: represents the linear effects of each factor x_i_, βii: represents the quadratic effects of each factor x_i_, ***ε***: is the experimental error.

The accuracy and reliability of the developed model were evaluated through a validation experiment performed under the predicted optimal conditions, and the relative errors between the experimental and predicted responses were determined.

### 4.6. Effect of chemical inducers on violacein production

The effect of three reported chemical inducers on violacein production was evaluated: tryptophan, formic acid [9], and ampicillin [17]. Stock solutions were freshly prepared and sterilized by filtration through a 0.22 μm pore-size membrane. Each inducer was tested at three concentrations within the synthetic medium formulation: 0.15–0.6 mg/mL for tryptophan; 0.08–0.32 mg/mL for formic acid and 6.25–25 μg/mL for ampicillin. Inducers were added prior to inoculation, and the medium pH was adjusted to 7.0. All treatments were carried out in triplicate, including an inducer-free control. Cultivations were conducted under the standard incubation parameters described above, but adapted to a working volume of 63 mL in 125 mL flasks. Following incubation, cells were harvested for subsequent analyses.

### 4.7. Effect of aeration and cultivation time

To evaluate the impacts of aeration and cultivation time on the growth and violacein production of *Janthinobacterium* sp. UV13, experiments were conducted in the optimized medium formulation supplemented with 0.3 mg/mL tryptophan (HPE-VP). To assess aeration, medium-to-flask volume ratios of 1:2 and 1:5 were compared. To assess cultivation time, cultures utilizing the 1:2 ratio were grown for either 72 or 144 h. All experimental treatments were performed in triplicate and maintained under the operational conditions described above. Volumetric violacein productivity (mg/L·h) was calculated for each tested condition.

### 4.8. Comparative Evaluation with Baseline Medium

To validate the efficiency of the optimized culture conditions, the HPE-VP medium operated under the highest-performing volume ratio and cultivation time, was compared against the baseline medium (3 g/L TSB supplemented with 3.6 g/L glucose) described by Alem *et al.* (2020) for *Janthinobacterium* sp. UV13.

On one side, the transcriptional activity of the violacein pathway under these different culture conditions was compared. Within the vioABCDE violacein biosynthetic gene cluster, *vio*A and *vio*D were selected to evaluate via RT-qPCR. Total RNA was extracted from bacterial cultures following an initial lysozyme treatment to facilitate cell lysis, followed by RNA purification using the RNeasy Mini Kit (QIAGEN, Germany), according to the manufacturer’s instructions. RNA samples were treated with DNase to remove residual genomic DNA, and cDNA was synthesized using random hexamer primers and M-MLV Reverse Transcriptase (New England Biolabs, USA). Relative gene expression was analyzed using the 2^-ΔCt^ method, with recA serving as the reference gene. Amplifications were performed using SensiFAST™ SYBR® No-ROX Kit (Bioline) on a DTprime Real-Time PCR System. Primer sequences and cycling conditions are provided in Supplementary Table S2.

Additionally, the downstream processing and pigment purification efficiency from both media were compared as well. At the 72 h endpoint, the culture broth underwent three sonication cycles (1 min per cycle; 30 s ON, 30 s OFF) in an ultrasonic probe sonicator (Vibra-Cell^TM^ VC750, Sonics^◻^) at 270 W, 20 Hz, and 40% amplitude. The resulting mixture was saturated with NaCL and extracted with n-butanol (6 mL per 25 mL of medium) under continuous agitation for 1 h. The pigmented butanol phase was washed with a saturated NaCL solution, recovered and dried until a constant weight. Samples were then sonicated for 20 min in 30 mL of a 1:1 H₂O:acetonitrile (MeCN) mixture. The solution was brought up to a final volume of 50 mL with the same solvent mixture; a 5 mL aliquot was then diluted to 10 mL, filtered and analyzed. Violacein-related compounds were identified by comparing their retention times, UV–Vis spectra, and MS/MS fragmentation patterns. For violacein quantitation in both samples, a calibration curve was prepared using a violacein standard.

The standard and the violacein extracts obtained from both medium conditions were analyzed via HPLC-DAD using a Shimadzu LC-2050C 3D liquid chromatograph. Chromatographic separation was performed on a Phenomenex Luna column (150 × 4.6 mm, 5 µm) at 30 °C, with a flow rate of 1.0 mL/min and a 10 µL injection volume. The mobile phase was 0.05% formic acid in water (A) and MeCN (B). Separation utilized a gradient from 10–98% B over 15 min; followed either by an immediate 3-min re-equilibration for standard (18 min total) or a 7-min column-cleaning plateau at 98%, prior to re-equilibration for extracts (25 min total). UV–Vis spectra were recorded at 190–800 nm, with violacein detection at 575 nm. LC–MS/MS analysis was performed on a Shimadzu LCMS-8040 system equipped with an SPD-M20A DAD detector and an electrospray ionization (ESI) source. The chromatographic separation matched the HPLC-DAD conditions, using a 1:4 flow split to the mass spectrometer. The MS was operated in full-scan mode (over the range of m/z 100–600) in both positive and negative ionization modes, with collision energies of ± 15, 25 and 35 V.

### 4.9. Statistical analysis

Descriptive statistics, including mean and standard deviation, were employed to examine the data. Additionally, two-dimensional graphs were utilized to facilitate data visualization. Experimental designs were performed using Protimiza Experimental Design Software. Statistical analyses were conducted by t-test and one-way ANOVA for multiple comparisons.The significance level employed was α = 0.05.

## Supporting information

Supplemental Pazos 2026

## 6. Author contributions

L.C. and D.A. designed the experiments, analyzed and interpreted the data, conducted scientific direction and wrote the manuscript.

H.P-E. designed and performed the experiments, acquired, analyzed and interpreted the data and wrote the manuscript.

E.V. designed, acquired, analyzed and interpreted the data of the bioprocess experiments

G.V and M.B. designed, acquired, analyzed and interpreted the data of the purification and structural determination experiments

A. C. performed the RT-qPCR experiments, acquired, analyzed and of the bioprocess experiments

L.C., D.A., H.P-E., E.V. G.V., M.B. and A.C. discussed results and provided manuscript feedback.

## 7 - Acknowledgements

The authors would like to thank Susana Castro-Sowinski (Biochemistry Section, Faculty of Sciences, Universidad de la República, Montevideo, Uruguay) for kindly providing the *Janthinobacterium* sp. UV13 strain used in this study.

## 8. Data availability statement

All data supporting the findings of this study are available within the article and its extended data files, or from the corresponding author upon reasonable request.

## References

1. Durán, N. et al. Multi-target drug with potential applications: violacein in the spotlight. World J. Microbiol. Biotechnol. 37, 151–151; 10.1007/s11274-021-03120-4 (2021).

2. Sedláček, I., et al. Classification of a Violacein-Producing Psychrophilic Group of Isolates Associated with Freshwater in Antarctica and Description of Rugamonas violacea sp. nov.; 10.1128/Spectrum (2021).

3. Choi, S. Y., Lim, S., Yoon, K. hye, Lee, J. I. & Mitchell, R. J. Biotechnological Activities and Applications of Bacterial Pigments Violacein and Prodigiosin. Journal of Biological Engineering. 15; 10.1186/s13036-021-00262-9 (2021).

4. Hoshino, T. Violacein and related tryptophan metabolites produced by Chromobacterium violaceum: biosynthetic mechanism and pathway for construction of violacein core. Appl. Microbiol. Biotechnol. 91, 1463–1475 (2011).

5. Ahmed, A. et al. Recent advances in synthetic, industrial and biological applications of violacein and its heterologous production. Journal of Microbiology and Biotechnology. 31, 1465–1480; 10.4014/jmb.2107.07045 (2021).

6. Durán, N. et al. New Advancements in Production, Purification, and Current Biological Activities of Violacein. Journal of the Brazilian Chemical Society. 36 10.21577/0103-5053.20250096 (2025).

7. Mojib, N., Farhoomand, A., Andersen, D. T. & Bej, A. K. UV and cold tolerance of a pigment-producing Antarctic Janthinobacterium sp. Ant5-2. Extremophiles 17, 367–378 (2013).

8. Xu, X. et al. Complete genome sequence of Janthinobacterium sp. B9-8, a violacein-producing bacterium isolated from low-temperature sewage. Microb. Pathog. 128, 178–183 (2019).

9. Cheng, K. C., et al. Improvement in Violacein Production by Utilizing Formic Acid to Induce Quorum Sensing in Chromobacterium violaceum. Antioxidants 11, (2022).

10. Gonçalves, P. R. et al. Violacein induces death of RAS-mutated metastatic melanoma by impairing autophagy process. Tumor Biology 37, 14049–14058 (2016).

11. Choi, S. Y., Im, H. & Mitchell, R. J. Violacein and bacterial predation: promising alternatives for priority multidrug resistant human pathogens. Future Microbiol. 12, 835–838 (2017).

12. Olivera-Bravo, S. et al. Neuroprotective effects of violacein in a model of inherited amyotrophic lateral sclerosis. Sci. Rep. 12, 4439 (2022).

13. Mendes, A. S., De Carvalho, J. E., Duarte, M. C. T., Durán, N. & Bruns, R. E. Factorial Design and Response Surface Optimization of Crude Violacein for Chromobacterium Violaceum Production. Biotechnology Letters. 23 (2001).

14. Wang, H. et al. Optimization of culture conditions for violacein production by a new strain of Duganella sp. B2. Biochem. Eng. J. 44, 119–124 (2009).

15. Fang, M. Y. et al. High crude violacein production from glucose by Escherichia coli engineered with interactive control of tryptophan pathway and violacein biosynthetic pathway. Microb. Cell Fact. 14, (2015).

16. Sun, H., Zhao, D., Xiong, B., Zhang, C. & Bi, C. Engineering Corynebacterium glutamicum for violacein hyper production. Microb. Cell Fact. 15, (2016).

17. Kanelli, M. et al. Microbial production of violacein and process optimization for dyeing polyamide fabrics with acquired antimicrobial properties. Front. Microbiol. 9, (2018).

18. Lu, Y. et al. Production of violet pigment by a newly isolated psychrotrophic bacterium from a glacier in Xinjiang, China. Biochem. Eng. J. 43, 135–141 (2009).

19. Atalah, J. et al. Isolation and characterization of violacein from an Antarctic Iodobacter: a non-pathogenic psychrotolerant microorganism. Extremophiles 24, 43–52 (2020).

20. Sigma-Aldrich. Violacein from Janthinobacterium lividum, >98% (HPLC) - V9389. https://www.sigmaaldrich.com/US/en/product/sigma/v9389. (2026).

21. Alem, D. et al. Production and antiproliferative effect of violacein, a purple pigment produced by an Antarctic bacterial isolate. World J. Microbiol. Biotechnol. 36, (2020).

22. Sasidharan, A. et al. Antifungal activity of violacein purified from a novel strain of Chromobacterium sp. NIIST (MTCC 5522). Journal of Microbiology 53, 694–701 (2015).

23. Kilonzo, P. M. & Margaritis, A. The effects of non-Newtonian fermentation broth viscosity and small bubble segregation on oxygen mass transfer in gas-lift bioreactors: A critical review. Biochem. Eng. J. 17, 27–40 (2004).

24. Schulze, C., et al. Investigation of exopolysaccharide formation and its impact on anaerobic succinate production with Vibrio natriegens. Microb Biotechnol 17 (1), 10.1111/1751-7915.14277 (2024).

25. Lee, M. E., Aswani, A., Han, A. S., Tomlin, C. J. & Dueber, J. E. Expression-level optimization of a multi-enzyme pathway in the absence of a high-throughput assay. Nucleic Acids Res. 41, 10668–10678 (2013).

26. Calemi, D. B. de A., et al. Evaluation of Violacein Metabolic Stability and Metabolite Identification in Human, Mouse, and Rat Liver Microsomes. Pharmaceutics 17, (2025).

27. Shuler, M., Kargi, F. & DeLisa, M. Bioprocess Engineering Basic Concepts (ed. Pearson, London, UK, 2017).

28. Reid, S. & Abratt, V. Sucrose utilisation in bacteria: genetic organisation and regulation. Appl. Microbiol. Biotechnol. 67, 312–321 (2005).

29. Gharat, K. & Singhal, R. A strategic media engineering and fed-batch approach enhances violacein production by Chromobacterium violaceum MTCC2656. Systems Microbiology and Biomanufacturing 4, 1106–1120 (2024).

30. Sutherland, I. The biofilm matrix–an immobilized but dynamic microbial environment. Trends Microbiol. 9, 222–227 (2001).

31. Zhang, Q. et al. A comprehensive review on violacein production by microbial fermentation. BioDesign Research. 7; 10.1016/j.bidere.2025.100043 (2025).

32. Frediansyah, A., Manuhara, Y. S. W., Kristanti, A. N., Luqman, A. & Wibowo, A. T. Fermentation in Minimal Media and Fungal Elicitation Enhance Violacein and Deoxyviolacein Production in Two Janthinobacterium Strains. Fermentation 8, (2022).

33. McClean, K. H. et al. Quorum sensing and Chrornobacteriurn violaceurn : exploitation of violacein production and inhibition for the detection of N-acyl homoserine lactones. Microbiology. 143, 3703–3740 (1997).

34. Ravichandran, V. et al. Virtual screening and biomolecular interactions of CviR-based quorum sensing inhibitors against Chromobacterium violaceum. Front. Cell. Infect. Microbiol. 8, 292–292 (2018).

35. Mion, S. et al. Disrupting quorum sensing alters social interactions in Chromobacterium violaceum. NPJ Biofilms Microbiomes 7, 40 (2021).

36. Bentley, R. & Haslam, E. The shikimate pathway—a metabolic tree with many branche. Crit. Rev. Biochem. Mol. Biol. 25, 307–384 (1990).

37. Akashi, H. & Gojobori, T. Metabolic Efficiency and Amino Acid Composition in the Proteomes of Escherichia Coli and Bacillus Subtilis. 99, (2002).

38. Yadav, V., De Mey, M., Lim, C., Ajikumar, P. & Stephanopoulos, G. The future of metabolic engineering and synthetic biology: Towards a systematic practice. Metab. Eng. 14, 233–241 (2012).

39. Bagoghli, N. & Hosseini-Abari, A. Improvement of violacein production using abiotic stresses and microbial adaptation. World J. Microbiol. Biotechnol. 40, (2024).

40. Balibar, C. & Walsh, C. In vitro biosynthesis of violacein from L-tryptophan by the enzymes VioA-E from Chromobacterium violaceum. Biochemistry 45, 15444–57 (2006).

41. Tong, Y., Zhou, J., Zhang, L. & Xu, P. A Golden-Gate Based Cloning Toolkit to Build Violacein Pathway Libraries in Yarrowia lipolytica. ACS Synth. Biol. 10, 115–124 (2021).

42. Choi, S., Kim, S., Lyuck, S., Kim, S. & Mitchell, R. High-level production of violacein by the newly isolated Duganella violaceinigra str. NI28 and its impact on Staphylococcus aureus. Sci. Rep. 5, 15598–15598 (2015).

43. Albalasmeh, A. A., Berhe, A. A. & Ghezzehei, T. A. A new method for rapid determination of carbohydrate and total carbon concentrations using UV spectrophotometry. Carbohydr. Polym. 97, 253–261 (2013).

