## Supplemental Pazos 2026 for "Enhancing violacein production in an Antarctic bacterial strain: New opportunities for biomedical use"

**Supplementary Table S1. Plackett–Burman design matrix for the eight components studied, including three center points**.

| **RUNS** | **X1** | **X2** | **X3** | **X4** | **X5** | **X6** | **X7** | **X8** |
| --- | --- | --- | --- | --- | --- | --- | --- | --- |
| **Flask_1** | 1 | -1 | -1 | 1 | -1 | -1 | 1 | 1 |
| **Flask_2** | 1 | 1 | -1 | -1 | 1 | -1 | -1 | 1 |
| **Flask_3** | -1 | 1 | 1 | 1 | -1 | -1 | -1 | -1 |
| **Flask_4** | 1 | -1 | -1 | 1 | 1 | 1 | -1 | -1 |
| **Flask_5** | 1 | 1 | 1 | -1 | 1 | -1 | 1 | -1 |
| **Flask_6** | 1 | 1 | 1 | 1 | -1 | 1 | -1 | 1 |
| **Flask_7** | -1 | 1 | -1 | 1 | 1 | 1 | 1 | -1 |
| **Flask_8** | -1 | -1 | 1 | 1 | 1 | -1 | 1 | 1 |
| **Flask_9** | -1 | -1 | 1 | -1 | 1 | 1 | -1 | 1 |
| **Flask_10** | 1 | -1 | 1 | -1 | -1 | 1 | 1 | -1 |
| **Flask_11** | -1 | 1 | -1 | -1 | -1 | 1 | 1 | 1 |
| **Flask_12** | -1 | -1 | -1 | -1 | -1 | -1 | -1 | -1 |
| **Flask_13** | 0 | 0 | 0 | 0 | 0 | 0 | 0 | 0 |
| **Flask_14** | 0 | 0 | 0 | 0 | 0 | 0 | 0 | 0 |
| **Flask_15** | 0 | 0 | 0 | 0 | 0 | 0 | 0 | 0 |

**Supplementary Table S2. Primer sequences and amplification conditions used for RT-qPCR analysis of vioA, vioD, and recA in *Janthinobacterium* sp. UV13.** Amplifications were performed using the SensiFAST™ SYBR® No-ROX Kit (Bioline) on a DTprime Real-Time PCR System, with an annealing temperature of 60 °C for 40 cycles.

| Gene | Primer | Sequence (5′–3′) | Amplicon size (bp) |
| --- | --- | --- | --- |
| vioA | vioA-F  vioA-R | CCTTTCACCCAGGCGGTATT  GTGGCTGACGAAATCCAGGA | 120 |
| vioD | vioD-F  vioD-R | ATCGACGGCCTGTTCAATCC  ACTTTCTGCTTGCGGCTTTG | 104 |
| recA | recA-F  recA-R | AAGCCCACTTCGACATCCTG  TAGCTGTACCAGGAACCGGA | 109 |

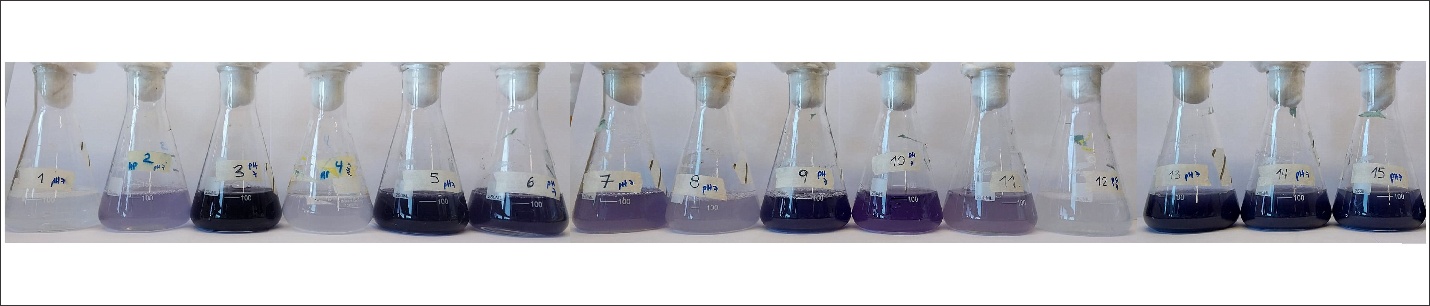
**Supplementary Figure S1. Screening of the synthetic medium components via a Plackett–Burman design.** Image shows photograph of culture flasks after 72 h of incubation under each tested condition. Quantitative data for these runs are provided in Table 1 of Section 2.2.

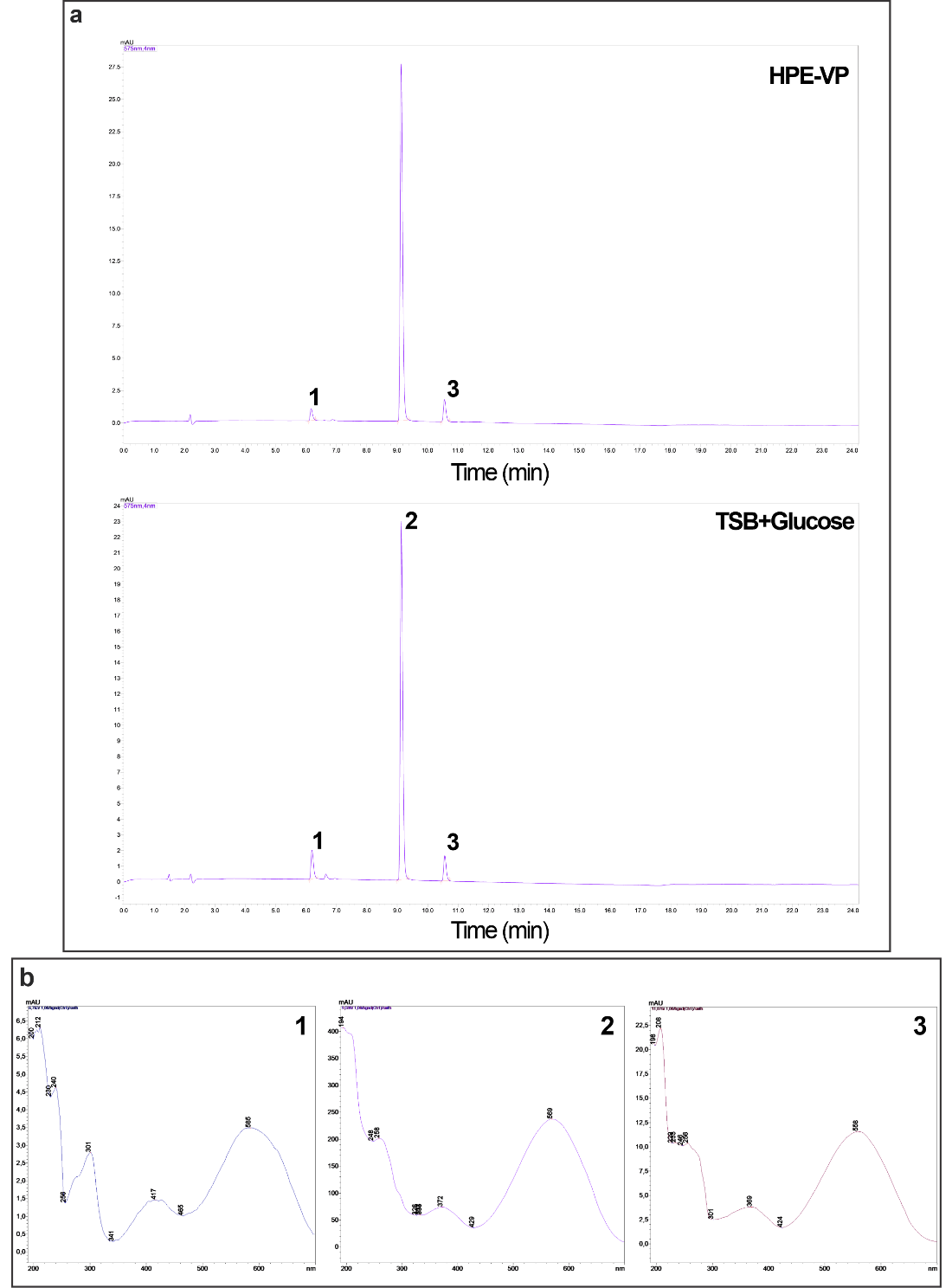

**Supplementary Figure S2. Identification and characterization of violacein-related compounds.** (a) Chromatographic profiles at 575 nm of the purified violacein extracts from HPE-VP and TSB+Glucose culture media. (b) UV–Vis absorbance spectra of the three peaks detected in each condition.

**LC–MS/MS analysis of the violacein extracts obtained from both medium conditions was performed as described in Section 4.8.** The MS was operated in full-scan mode (over the range of m/z 100–600) in both positive and negative ionization modes, with collision energies of ± 15, 25 and 35 V.

**Violacein (Peak 2, Retention time 9.1 min)**

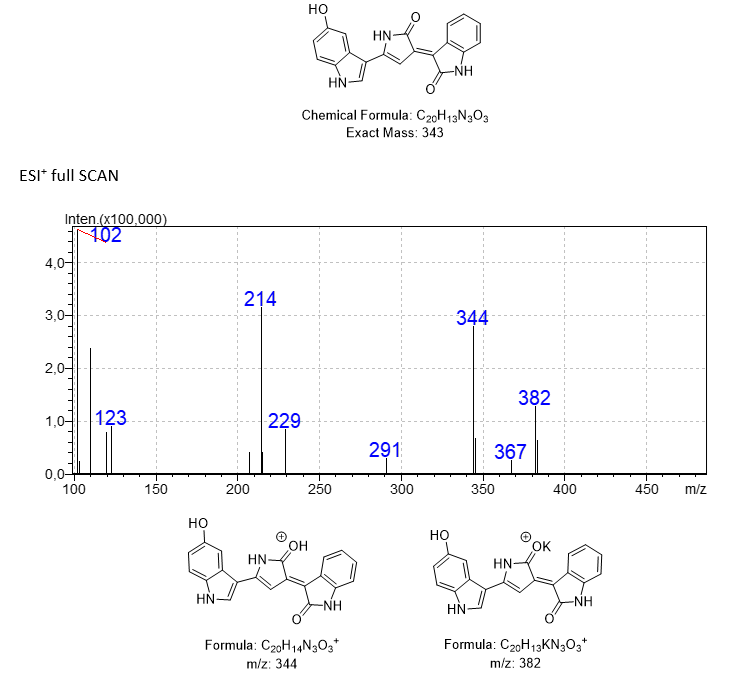

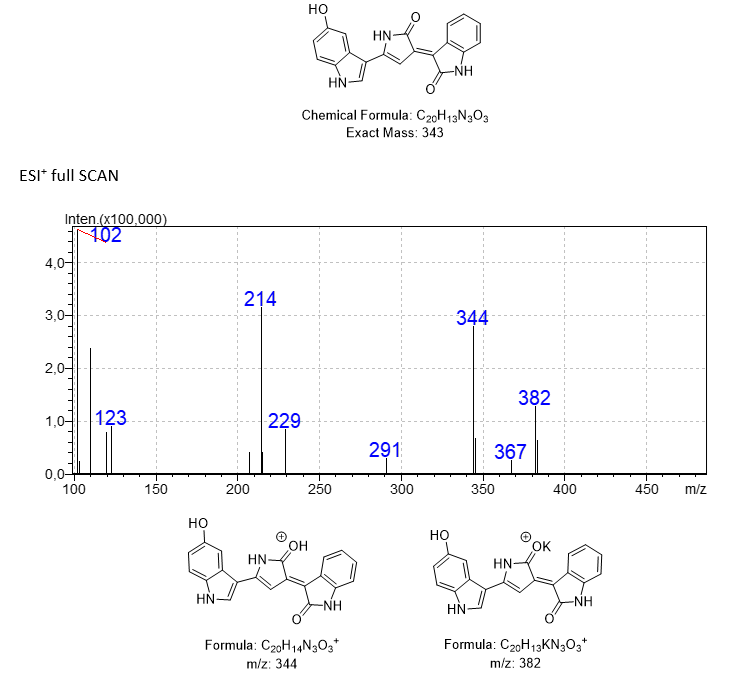

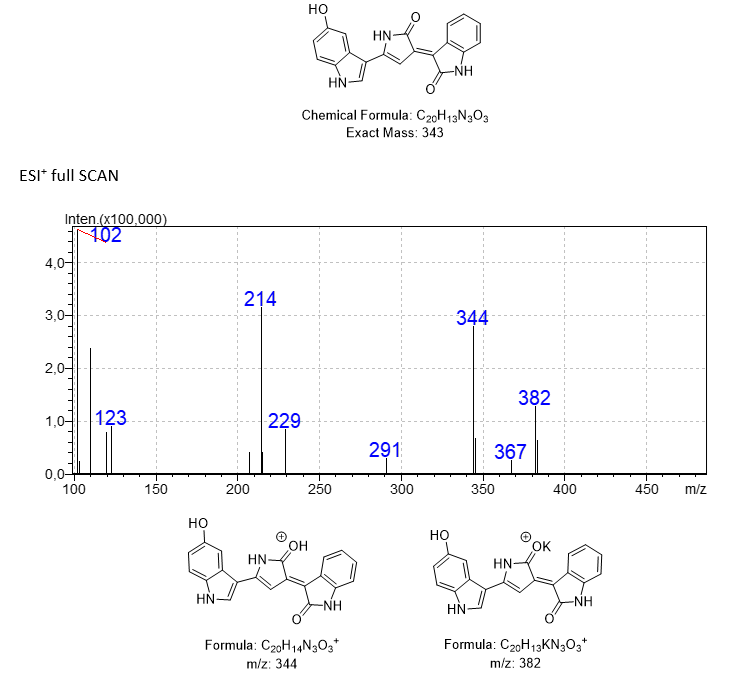

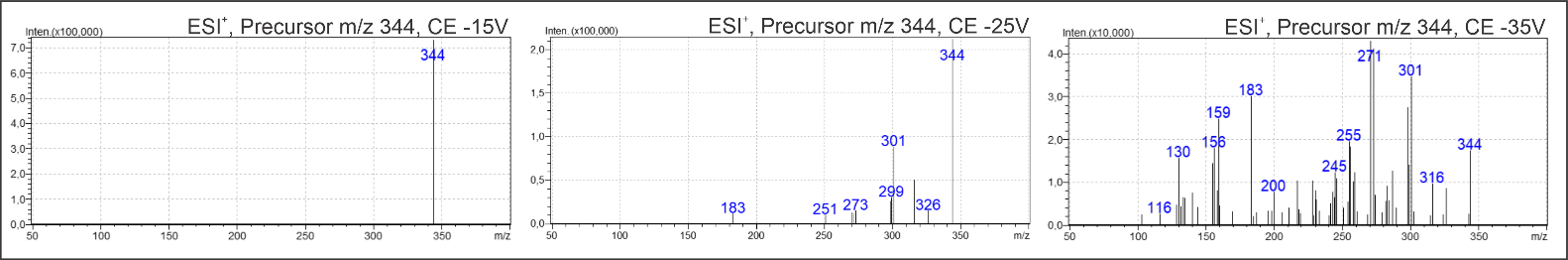

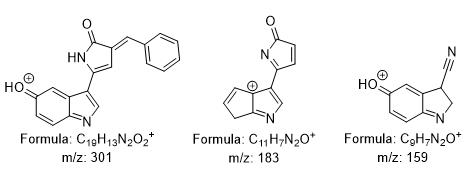
Proposed fragments according to Calemi *et al*. [25]

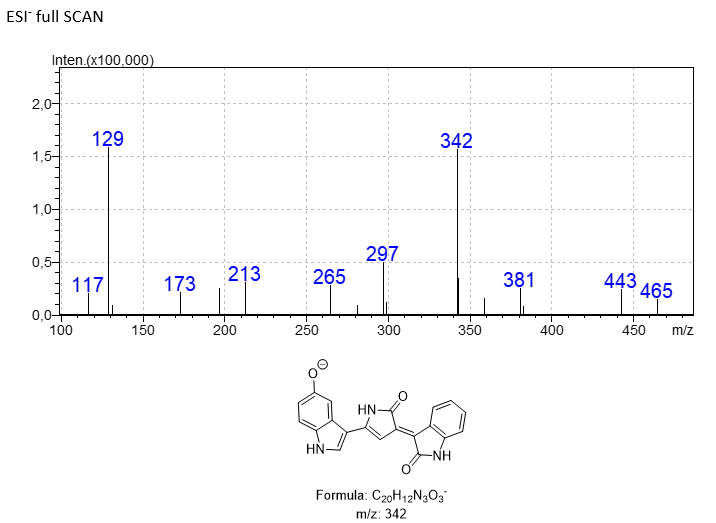

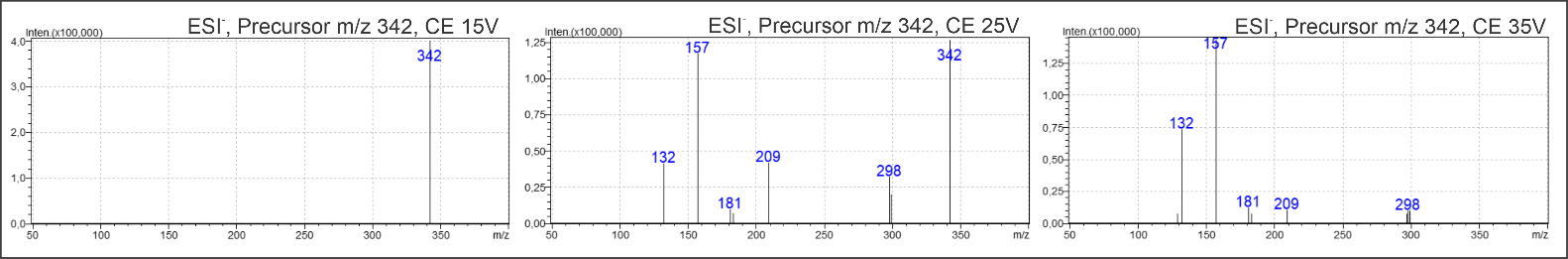

Proposed fragments according to Calemi *et al*. [25]
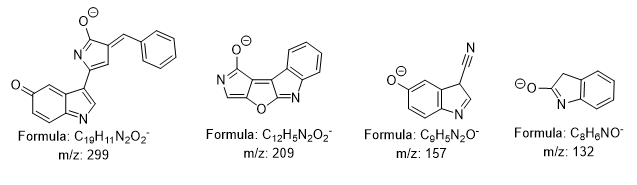

**Deoxyviolacein (Peak 3, Retention time 10.6 min)**

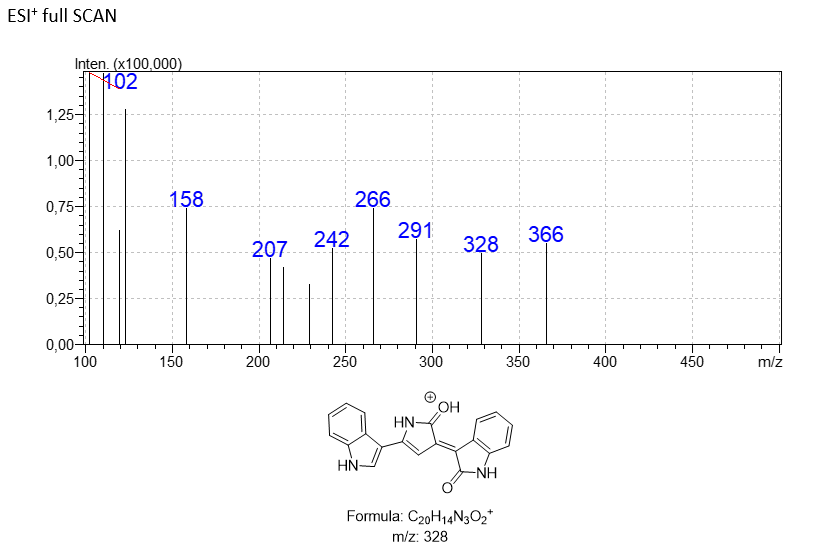

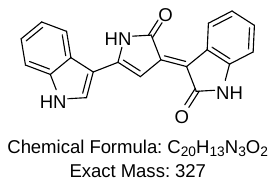

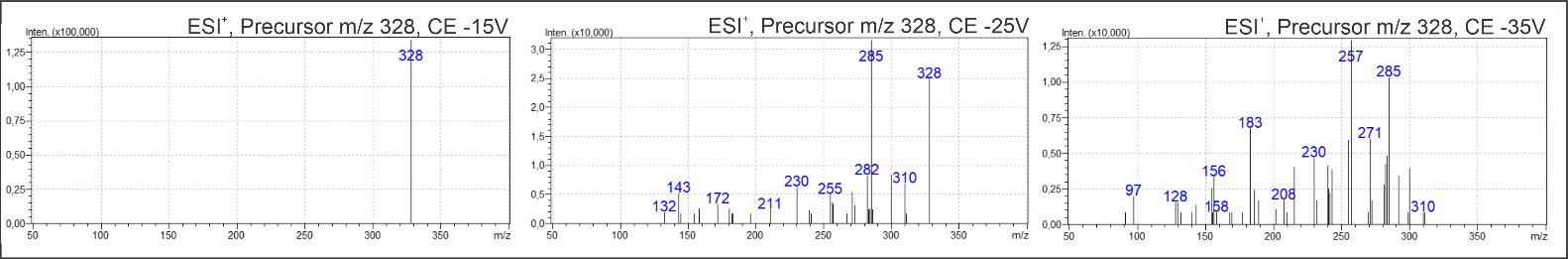

Proposed fragments according to Calemi *et al*. [25]

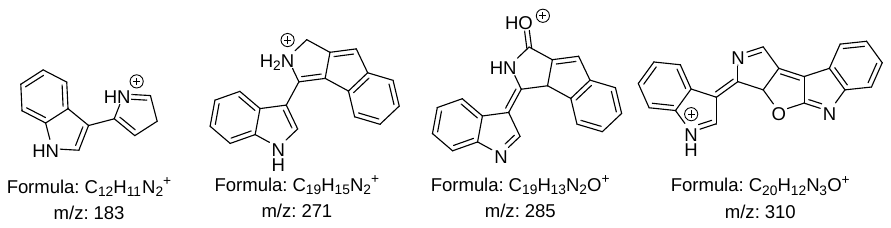

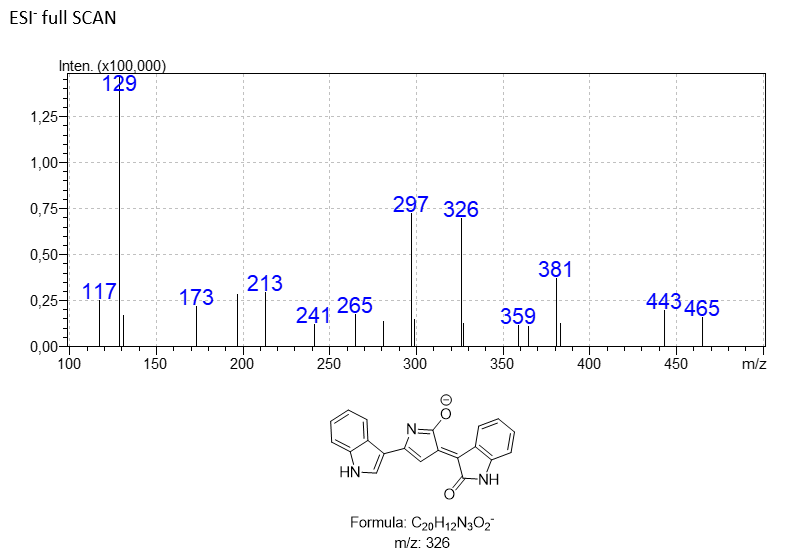

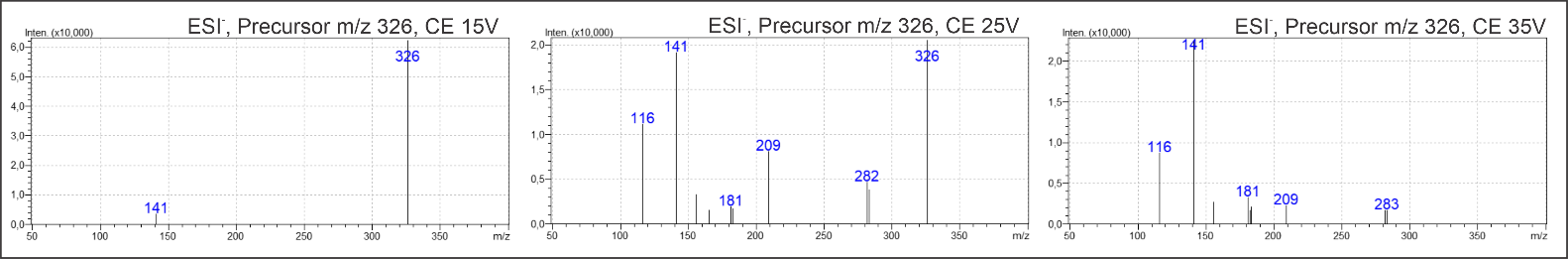

Proposed fragments according to Calemi *et al*. [25]

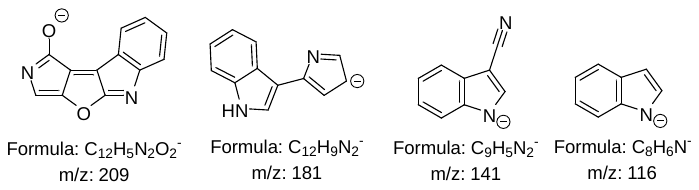

**Proviolacein (Peak 1, Retention time 6.2 min)**

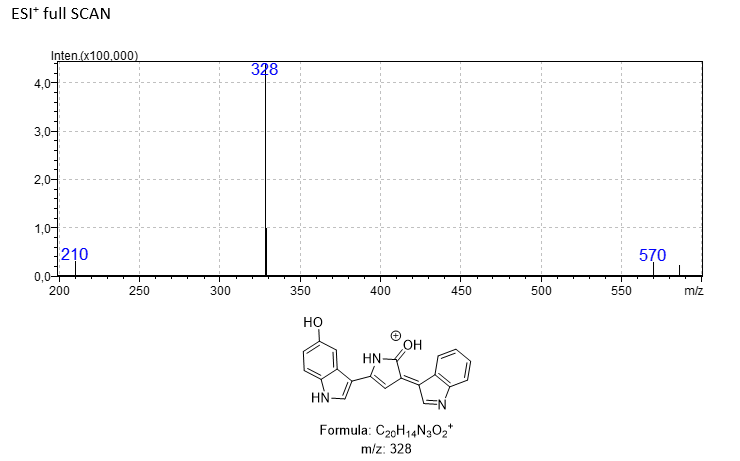

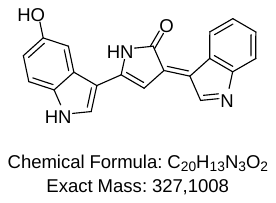

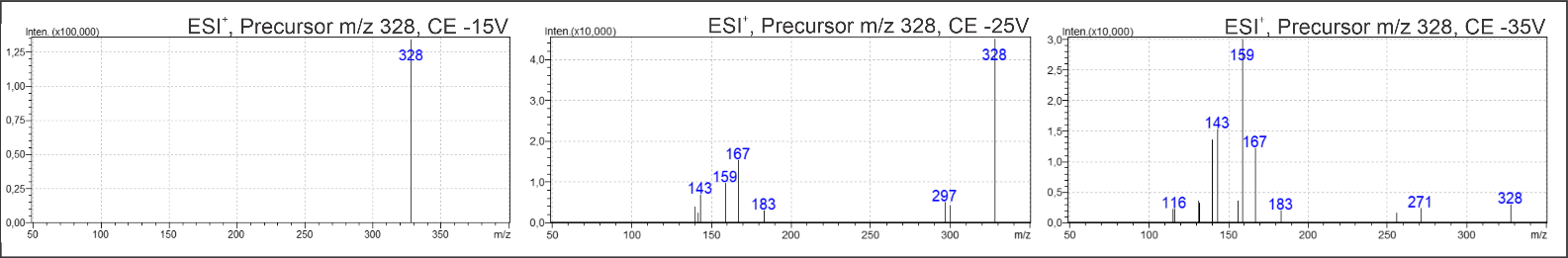

Proposed fragments according to Calemi *et al*. [25]

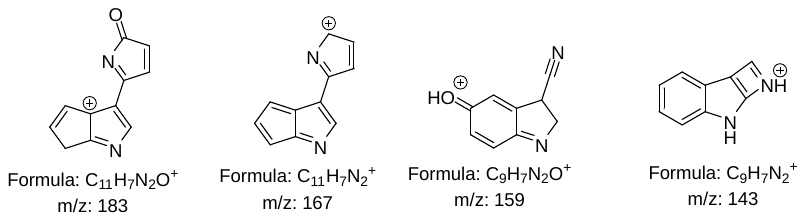

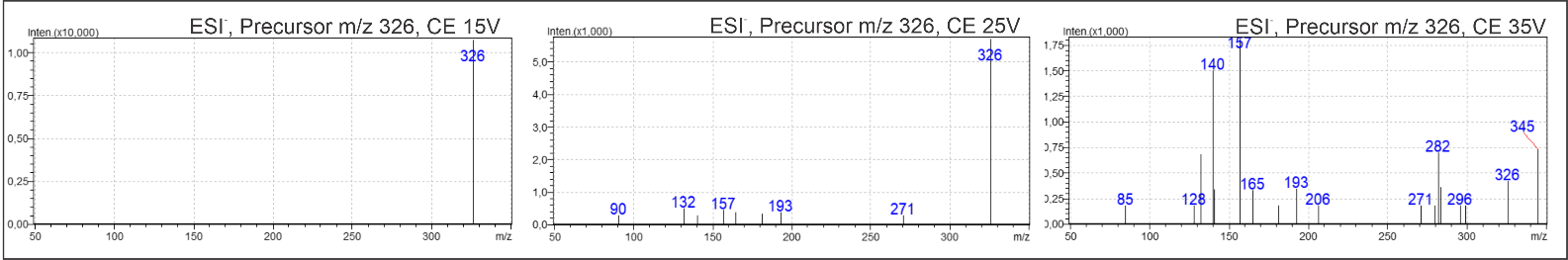

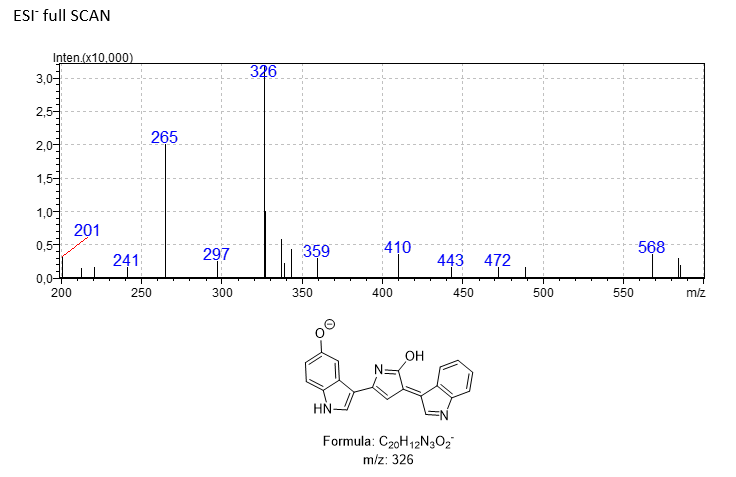

Proposed fragments according to Calemi *et al*. [25]

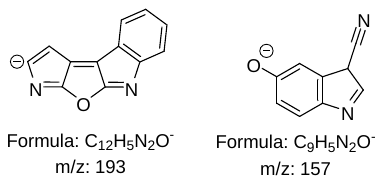
